# Ptbp1 deletion does not induce glia-to-neuron conversion in adult mouse retina and brain

**DOI:** 10.1101/2021.10.04.462784

**Authors:** Thanh Hoang, Dong Won Kim, Haley Appel, Nicole A. Pannullo, Patrick Leavey, Manabu Ozawa, Sika Zheng, Minzhong Yu, Neal S. Peachey, Juhyun Kim, Seth Blackshaw

## Abstract

Direct reprogramming of glia into neurons is a potentially promising approach for the replacement of neurons lost to injury or neurodegenerative disorders. Knockdown of the polypyrimidine tract-binding protein *Ptbp1* has been recently reported to induce efficient conversion of retinal Müller glia and brain astrocytes into functional neurons. However, genetic analysis of *Ptbp1* function in adult glia has not been conducted. Here, we use a combination of genetic lineage tracing, scRNA-Seq, and electrophysiological analysis to show that specific deletion of *Ptbp1* in adult retinal Müller glia and brain astrocytes does not lead to any detectable level of glia-to-neuron conversion. Only a few changes in gene expression are observed in glia following *Ptbp1* deletion, and glial identity is maintained. These findings highlight the importance of using genetic manipulation and lineage tracing methods in studying cell type conversion.

## Introduction

Neurodegenerative disorders are clinically diverse and represent a huge public health burden. To address this, considerable effort has been focused on directed transdifferentiation of endogenous glial cells into functional neurons that were lost to diseases. Most strategies to achieve this have included glial-specific viral overexpression of genes that promote neuronal identity, including transcription factors and miRNAs^1–6^, or genetic or small molecule-based manipulation of extracellular signaling pathways^7,8^. These approaches have achieved variable rates of success, with many treatments working only *in vitro* or in immature glia, inducing only partial reprogramming, or not generating desired neuronal subtypes^9^. Furthermore, studies that reported highly efficient reprogramming using these approaches have been criticized for lacking data that unambiguously demonstrate a lineage relationship between reprogrammed glia and neurons^10,11^.

The ability to reliably and efficiently induce glial reprogramming *in vivo* by manipulation of a single factor would be a major advance towards effective cell-based therapy for neurodegenerative disorders. Several recent reports have claimed to achieve exactly this outcome through knockdown of *Ptbp1* expression in retinal Müller glia and brain astrocytes^12–15^. *Ptbp1* is an RNA binding protein and splicing regulator that is broadly expressed in non-neuronal and neuronal progenitor cells and represses neuronal-specific alternative splicing^16–18^. Neural progenitor-specific deletion of *Ptbp1* leads to precocious neurogenesis^19^, and knockdown of *Ptbp1* has been reported to be sufficient to convert both fibroblast and N2a cells into neurons *in vitro*^*20*^. Furthermore, several recent papers reported that knockdown of *Ptbp1* is sufficient to induce glial conversion into neurons. One study reported that *Ptbp1* knockdown in retinal Müller glia using AAV-mediated CasRx led to rapid and efficient transdifferentiation of Müller glia into retinal ganglion cells, which were then able to efficiently innervate targets in the brain following excitotoxic inner retinal injury^12^. This same study also reported that *Ptbp1* knockdown was sufficient for efficient conversion of brain astrocytes into dopaminergic neurons in the striatum and rescued function in a 6-OHDA-induced mouse model of Parkinson’s disease (PD). A second study showed that lentiviral-mediated shRNA and antisense oligonucleotides (ASO)-mediated knockdown of *Ptbp1* in astrocytes in the cortex, striatum, and substantia nigra all induced efficient reprogramming of astrocytes into functional neurons, which in turn also rescued neurological defects in this same PD model^13^. Other studies have reported that *Ptbp1* knockdown can convert retinal Müller glia to photoreceptors^15^, and restore neurogenic competence in neural progenitor cells in the dentate gyrus of aged mice^14^.

This simple and elegant loss of function approach potentially addresses many of the difficulties associated with previous efforts towards directed glial reprogramming, particularly in a clinical setting, such as low reprogramming efficiencies and the use of complex overexpression constructs. However, several major concerns remain to be addressed. First, none of these approaches convincingly demonstrated a reduction of *Ptbp1* expression in glial cells *in situ*. Second, lineage relationships between glia and neurons were inferred through the use of GFAP promoter-based AAV constructs or transgene, which are known to show neuronal expression in some contexts^21,22^. Third, convincing evidence for direct glia-to-neuron conversion using reliable genetic lineage analysis and/or scRNA-Seq-based trajectory analysis is lacking. These concerns need to be addressed before the *Ptbp1* knockdown approach can further advance toward clinical applications.

In this study, we address the question of knockdown specificity and efficiency of glia-to-neuron conversion upon *Ptbp1* reduction, through the use of glial-specific conditional mutants of *Ptbp1*. We combined both genetic lineage and scRNA-Seq analysis of adult wildtype Müller glia and astrocytes, as well as adult glia carrying heterozygous or homozygous mutants of *Ptbp1*. Although we observe efficient and cell-specific disruption of *Ptbp1*, we observe no evidence for conversion of Müller glia or astrocytes into neurons in either heterozygous or homozygous *Ptbp1* mutants. ScRNA-Seq analysis reveals only subtle changes in gene expression in mutant glia, but no evidence for induction of neuronal-specific genes and no presence of neuronal physiology in mutant glia. Our data indicate that the glia-to-neuron conversion reported in previous studies following *Ptbp1* knockdown does not reflect the effects of *Ptbp1* loss of function.

## Results

### Genetic loss of function of *Ptbp1* in retinal Müller glia did not lead to glia-to-neuron conversion

To simultaneously disrupt *Ptbp1* in adult retinal Müller glia and irreversibly label these cells with a visible marker, we used three transgenic lines: *Glast*^*CreERT2*^, which efficiently and selectively induces Cre-dependent recombination in Müller glia following tamoxifen treatment^23,24^; *Sun1-GFP*^*lox/lox*^, which expresses GFP targeted to the nuclear envelope under the ubiquitous CAG promoter following Cre activation^25^; and *Ptbp1*^*lox/lox*^, in which *loxP* sites flank the promoter and the first coding exon of *Ptbp1*, and Cre activation disrupts transcription^19^. We then generated wildtype (*Glast*^*CreERT2*^;*Sun1-GFP*^*lox/lox*^; *Ptbp1*^*+/+*^), heterozygous (*Glast*^*CreERT2*^;*Sun1-GFP*^*lox/lox*^; *Ptbp1*^*lox/+*^), and homozygous (*Glast*^*CreERT2*^;*Sun1-GFP*^*lox/lox*^; *Ptbp1*^*lox/lox*^) mutant mice. Starting at ∼5 weeks old, we induced Cre activation by using 4 doses of daily injections of 4-Hydroxytamoxifen (4-OHT) and conducted immunohistochemical analysis 2 and 4 weeks later (Fig.1a).

**Figure 1.**
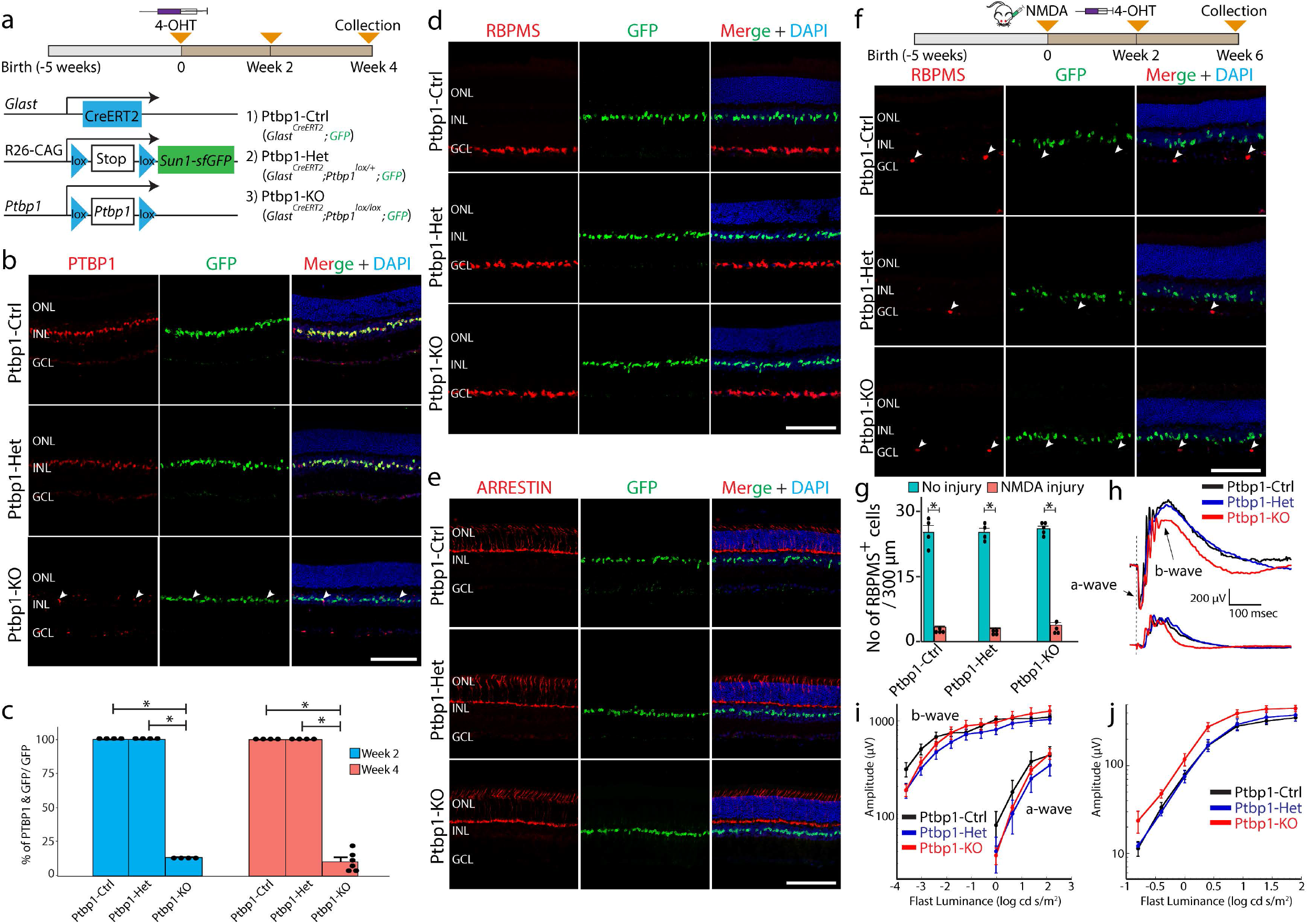
*Ptbp1* deletion does not result in the conversion of Müller glia into retinal neurons in adult mice. **a**, A schematic diagram of the generation of specific deletion of *Ptbp1* and lineage tracing of Müller glia. **b**, Representative immunohistochemistry of PTBP1 expression in wildtype (Ptbp1-Ctrl), heterozygous (Ptbp1-Het), and homozygous (Ptbp1-KO) *Ptbp1* retinas after 4 weeks of 4-OHT i.p injection. White arrowheads indicate residual Ptbp1-expressing Müller glial cells. **c**, Quantification of the percentage of PTBP1/GFP-double positive Müller glial cells after 2 and 4 weeks of 4-OHT i.p injection (n >= 4 retinas/genotype). **d**, Representative immunohistochemistry for RBPMS, a pan-retinal ganglion cell marker, in retinal sections from three genotypes (n >= 6 retinas/genotype) after 4 weeks of 4-OHT i.p injection. **e**, Representative images of immunohistochemical analysis of cone arrestin in retinal sections from three genotypes after 4 weeks of 4-OHT i.p injection (n >= 3 retinas/genotype). **f**, Representative immunohistostaining of RBPMS in retinal sections from the three genotype groups at 6 weeks after NMDA injury (n >= 3 retinas/genotype). White arrowheads indicate residual RBPMS-expressing ganglion cells. **g**, Quantification of RBPMS-positive cells in wildtype, heterozygous, and homozygous *Ptbp1* mutant retinas (n >= 4 retinas/genotype) with and without NMDA injury. **h**, Representative waveforms for dark and light-adapted ERGs. No obvious defect is seen in *Ptbp1*-deficient retinas. **i**, Dark-adapted and **j**. light-adapted ERGs as a function of stimulus intensity (n = 5 mice/genotype). No differences are seen between *Ptbp1*-deficient retinas and controls *(p > 0*.*06*). ONL, outer nuclear layer; INL, inner nuclear layer; GCL, ganglion cell layer. Scale bars = 100 µm.

Immunohistochemistry data showed that PTBP1 protein expression is enriched in Müller glia and some other non-glial cells in adult wildtype retinas (Fig.1b). We observed that Cre activation led to a ∼ 90% reduction in the number of PTBP1-positive Müller glia cells at both 2 and 4 weeks following 4-OHT induction in the homozygous retinas (Fig.1b,c). GFP-positive nuclei of Müller glia remained confined to the inner nuclear layer, implying that they did not generate either retinal ganglion cells or photoreceptors (Fig.1b). To confirm this finding, we then conducted immunostaining for the retinal ganglion cell-specific markers RBPMS (Fig.1d, Extended Data Fig.1a, c) and BRN3B (Extended Data Fig.1d, e), in contrast to a recent report^12^, we did not observe any colocalization of these markers with GFP following *Ptbp1* deletion. We likewise observed no colocalization of the cone-specific marker arrestin (Fig.1e) or the photoreceptor and bipolar marker OTX2 (Extended Data Fig.1f) with GFP, in contrast to another recent study^15^. No induction of GFAP expression was observed in *Ptbp1*-deficient Müller glia (Extended Data Fig.1g), indicating that *Ptbp1* depletion did not initiate reactive gliosis. It had also been previously reported the robust conversion of Müller glia to retinal ganglion cells induced by *Ptbp1* depletion resulted in the restoration of visual function following excitotoxic NMDA damage^12^. Although NMDA treatment led to extensive loss of RBPMS-positive retinal ganglion cells, we likewise did not observe any GFP/RBPMS or GFP/BRN3B-positive cells nor the recovery of retinal ganglion cells after 4 weeks following NMDA injury (Fig.1f,g, Extended Data Fig.1b,h). ERG analysis did not reveal any obvious defects or significant differences between control and *Ptbp1*-deficient retinas in a-or b-wave amplitude dark-adapted animals or in b-wave amplitude in light-adapted animals (Fig. 1h-j). Taken together, our data demonstrate that loss of *Ptbp1* does not convert Müller glia into retinal neurons either in healthy or damaged retinas.

### Genetic loss of function of *Ptbp1* in brain astrocytes did not lead to glia-to-neuron conversion in the brain

To genetically disrupt *Ptbp1* function in astrocytes, we employed a similar approach as in the retina, using astrocyte-specific tamoxifen-inducible *Aldh1l1*^*CreERT2*^ mice^26^, as well as the S*un1-GFP*^*lox/lox*^ and *Ptbp1*^*lox/lox*^ lines. We used the same breeding and tamoxifen induction strategy as in the retina to generate mice that were wildtype, heterozygous and homozygous for *Ptbp1* loss of function in astrocytes, and at the same time, we irreversibly labeled astrocytes with Sun1-GFP for lineage tracing. Immunohistochemical analysis was then performed at 2 and 8 weeks following 4-OHT intraperitoneal (i.p) induction (Fig.2a).

**Figure 2.**
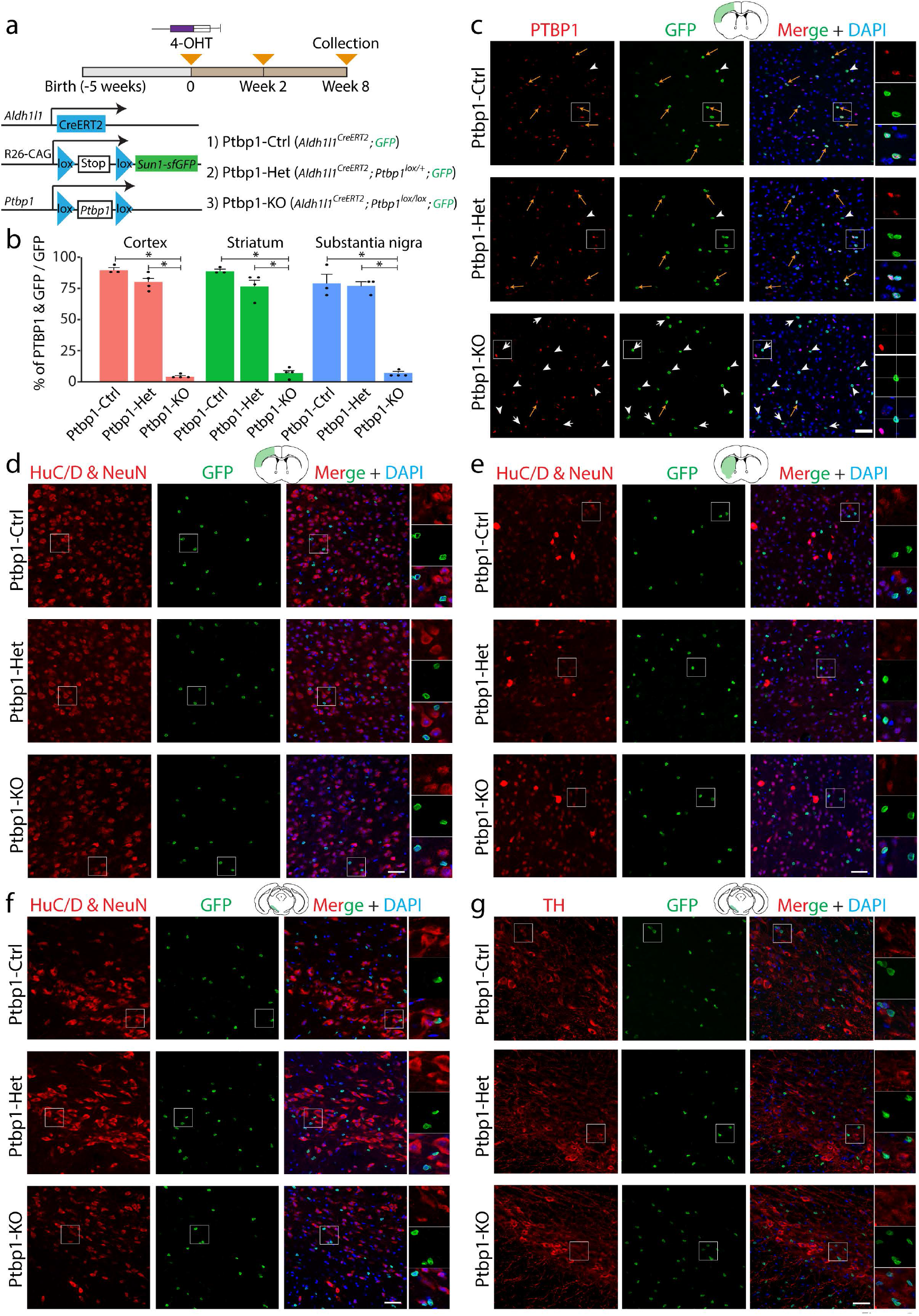
Loss of Ptbp1 did not result in astrocyte-to-neuron conversion in the adult mouse brain. **a**, A schematic diagram for the generation of specific deletion of *Ptbp1* and lineage tracing of astrocytes, and experimental timelines. **b**, Quantification of the percentage of PTBP1/GFP-double positive cells showing a highly efficient *Ptbp1* deletion in all three brain regions (cortex, striatum, and substantia nigra). **c**, Representative PTBP1 and GFP immunostaining images of the cortex at 8 weeks after 4-OHT i.p injection (n >= 3 mice/genotype). Yellow arrows indicate PTBP1/GFP-double positive cells. White arrowheads indicate PTBP1-negative/GFP-positive cells. **d-f**, Representative immunostaining images of HuC/D;NeuN and GFP in the cortex (**d**), striatum (**e**), and substantia nigra (**f)** at 8 weeks after 4-OHT i.p injection (n >= 3 mice/genotype). No HuCD;NeuN/GFP-double positive cells were observed in any of the three brain regions across all 3 genotypes. **g**, Representative immunostaining images of TH and GFP in the substantia nigra at 8 weeks after 4-OHT i.p injection. No TH/GFP-double positive cells were observed in the substantia nigra in all 3 genotypes. (n >=3 mice/genotype). Scale bars = 50 µm.

As previous studies have reported glia-to-neuron conversion following *Ptbp1* knockdown in cortex, striatum, and substantia nigra^12,13^, we focused our analysis on these 3 brain regions. We observed a dramatic reduction in the percentage of PTBP1/GFP-double positive cells relative to total GFP-positive cells in homozygous mice in all three regions compared to wildtype, and a modest reduction in the heterozygotes relative to wildtype mice in cortex and striatum (Fig.2b,c, Extended Data Fig.2). However, we did not observe any GFP-positive cells co-labeled with the neuronal markers NeuN and HuC/D in these regions in any genotypes, at either 2 or 8 weeks following 4-OHT i.p induction (Fig.2d-f, Extended Data Fig.3a-c). Since previous studies had reported that *Ptbp1* knockdown induced markers of dopaminergic neurons in striatum and substantia nigra^12,13^, we then tested whether *Ptbp1* deletion induced either tyrosine hydroxylase (TH) or the dopaminergic transporter (DAT). However, we did not observe any colocalization of GFP with either marker in either region, regardless of the genotype (Fig. 2g, Extended Data Fig.3d-f).

We next examined if there is any potential effect of *Ptbp1* deletion on astrocyte physiology. No GFP-positive astrocytes in the cortex, in either *Ptbp1* heterozygotes and homozygotes, fired action potentials in response to the current steps in whole-cell patch-clamp recordings, whereas neighboring GFP-negative wildtype neurons readily fired action potentials (Fig.3a). This indicates that *Ptbp1* deletion did not change the electrophysiological functions of astrocytes to make them fire action potentials like neurons. The astrocytes displayed highly negative resting membrane potentials (Ptbp1-Het, -82.2 ± 1.1 mV; Ptbp1-KO, -81.4 ± 1.9 mV) as previously reported^27^, and the *Ptbp1* heterozygous and homozygous cells did not show a significant difference in resting membrane potential (Fig.3b, p=0.7795, Mann-Whitney test). These results suggest that *Ptbp1* deletion in astrocytes does not induce any specific neuronal-like electrophysiological changes.

**Figure 3.**
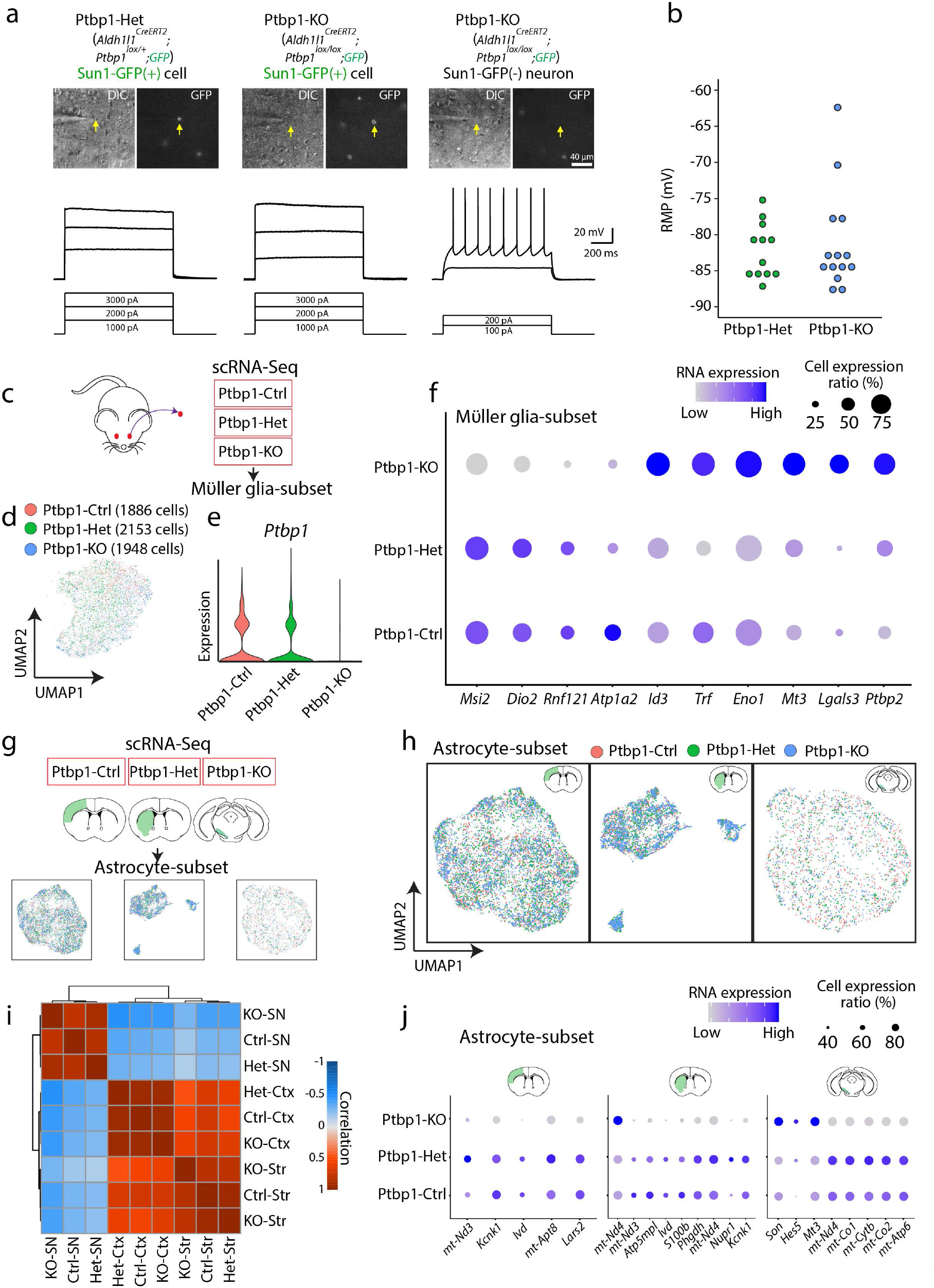
No changes in electrophysiological properties of cortical astrocytes and only subtle changes in gene expression in Müller glia and astrocytes are observed after *Ptbp1* deletion. **a**, Representative cell images and voltage traces to the depolarizing current steps of a GFP-positive cell from Ptbp1-Het (left), Ptbp1-KO mice (middle), and a GFP-negative neuron from a Ptbp1-KO mouse (right). Yellow arrows indicate the location of the recorded cells in bright and fluorescence images. Scale bar = 40 µm. **b**, A summary graph for resting membrane potential (RMP) of the GFP-positive cells between the two groups (Ptbp1-Het, 12 cells from 3 mice; Ptbp1-KO, 14 cells from 3 mice). **c**, A schematic diagram of scRNA-Seq experiments in *Glast*^*CreERT2*^;*Sun1-GFP*^*lox/lox*^; *Ptbp1*^*+/+*^ (Ptbp1-Ctrl), *Glast*^*CreERT2*^;*Sun1-GFP*^*lox/lox*^; *Ptbp1*^*lox/+*^ (Ptbp1-Het), *Glast*^*CreERT2*^;*Sun1-GFP*^*lox/lox*^; *Ptbp1*^*lox/lox*^ (Ptbp1-KO) retina. **d**, UMAP plot showing subsetted retinal Müller glia from all 3 genotypes. Note that there is no separated *Ptbp1* knockout Müller glia cluster(s). **e**, Violin plots showing *Ptbp1* RNA expression in Müller glia across the 3 genotypes. **f**, Dot plots showing some mildly changed gene expression of Müller glia across the 3 genotypes. **g**, A schematic diagram of scRNA-Seq experiments in *Aldh1l1*^*CreERT2*^;*Sun1-GFP*^*lox/lox*^; *Ptbp1*^*+/+*^ (Ptbp1-Ctrl), *Aldh1l1*^*CreERT2*^;*Sun1-GFP*^*lox/lox*^; *Ptbp1*^*lox/+*^ (Ptbp1-Het), *Aldh1l1*^*CreERT2*^;*Sun1-GFP*^*lox/lox*^; *Ptbp1*^*lox/lox*^ (Ptbp1-KO) in the brain. **h**, UMAP plot showing subsetted astrocytes in the cortex (left), striatum (middle) and substantia nigra (right) of all 3 genotypes. Note that there is no separated *Ptbp1* knockout astrocyte cluster(s). **i**, Heatmap showing correlations of subsetted astrocytes across genotypes and brain regions. **j**, Dot plots showing some mildly changed gene expression of subsetted astrocytes in the cortex (left), striatum (middle), and substantia nigra (right) across genotypes. Ctx = Cortex, Str = Striatum, SN = Substantia nigra.

### Genetic loss of function of *Ptbp1* leads to only subtle changes in the gene expression profile of Müller glia and astrocytes

Previous studies did not characterize the gene expression profile of glial cells after *Ptbp1* knockdown^12,13^. To comprehensively profile the cellular phenotype induced by *Ptbp1* deletion in Müller glia and astrocytes, we performed scRNA-Seq analysis of retina, cortex, striatum, and substantia nigra 2 weeks following tamoxifen treatment of wildtype, heterozygous, and homozygous mice (Fig.3c,g). Müller glia and astrocytes were then subsetted for further analysis (Fig.3d,h, Extended Data Fig.4a,b, Extended Data Fig.5a-c).

In all tissues, we see a consistent reduction in *Ptbp1* expression levels in Müller glia or astrocytes in heterozygous and homozygous mice (Fig.3e, Extended Data Fig.5d). The relative fraction of either Müller glia or any major subtype of retinal neurons did not change among any of the genotypes (Extended Data Fig.4b). We observed few significant changes in gene expression in any profiled glial population. In Müller glia, we observed increased expression of the *Ptbp1* paralogue *Ptbp2* expression level following *Ptbp1* disruption (Fig.3f). *Ptbp2* upregulation has previously been reported in neural progenitors following *Ptbp1* loss of function^16^, and since *Ptbp1* and *Ptbp2* show partially redundant functions^28^, this may provide some functional compensation for *Ptbp1* loss of function. We also observed increased expression of *Id3, Trf, Eno1, Mt3, Lgals3*, with reduced expression of *Msi2, Dio2, Rnf121, Atp1a2* (Fig.3f). In all cases, the changes in gene expression were quite modest (Table ST1); we did not observe either significant reductions of expression of Müller glia-specific markers, including *Sox9, Glul, Rlbp1, Slc1a3, Apoe, Aqp4, Mlc1, Kcnj10*, and *Tcf7l2*, or induction of genes specific to neural progenitors or mature neurons (Extended Data Fig.4d, Table ST1). Immunostaining confirmed that *Ptbp1*-deficient Müller glia retained expression of the glial marker SOX9 (Extended Data Fig.4e). Together, our data demonstrate that loss of function of *Ptbp1* does not dramatically alter gene expression in Müller glia, or cause these cells to lose their identity.

We likewise did not observe any clear changes in the astrocytes across the 3 genotypes in any of the brain regions (Fig.3h-i, Table ST2-ST4). A correlation plot showed that differences in gene expression in astrocytes among brain regions (cortex and striatum to substantia nigra) were higher than the differences among genotypes (Fig.3i). We also observed *S100b*-expressing astrocytes in the striatum as previously reported^29^ (Extended Data Fig.5e). As with Müller glia, we observed only very modest changes in gene expression following *Ptbp1* loss of function (Fig.3j, Table ST2-ST4). Upregulated genes include *mt-Nd4, Son, Hes5, Mt3*; downregulated genes include *mt-Nd3, Lars2, Ivd*. Most astrocyte markers did not show altered expression after *Ptbp1* deletion (Extended Data Fig.5f). Immunostaining confirmed that *Ptbp1*-deficient, GFP-positive astrocytes retained expression of glial marker SOX9 (Extended Data Fig.5g). We also did not observe induction of either neural progenitor or mature neuron-specific genes (Table ST2-ST4). We conclude that *Ptbp1* does not function in mature glial cells to repress the expression of neuronal genes.

## Discussion

Using the genetic loss of function and cell lineage analysis, in combination with scRNA-Seq analysis, we observed no evidence that either partial or complete loss of function of *Ptbp1* induces glia-to-neuron conversion in the retina or brain. Our data contrasts sharply with several recent studies that analyzed the effects of *Ptbp1* knockdown using ASO, shRNA, and/or CasRx^12–15^, but is in agreement with a recent study the re-examined previous studies reporting astrocyte-to-neuron conversion following *Neurod1* overexpression or either shRNA-or CasRx-mediated *Ptbp1* knockdown^30^. This latter study concluded that reports of glia-to-neuron conversion in these two models represented a leaky neuronal expression of the GFAP-based AAV and transgene lines used to label astrocytes. Leaky neuronal expression of GFAP-based reagents has been previously reported in other contexts^21,22^, and caution should be used when interpreting results obtained using these methods without corroborating data obtained using more strongly glial-specific minipromoters and Cre lines, such as the extensively validated *Glast*^*CreERT2*^ and *Aldh1l1*^*CreERT2*^ lines used here.

Our data show that the previous reports of glia-to-neuron conversion are unlikely to have resulted from *Ptbp1* loss of function in glial cells. Then what could have accounted for the recovery of visual function following NMDA excitotoxicity, or behavioral recovery following 6-OHDA-induced PD^12,13^? Ectopic neuronal expression of GFAP-based reagents in native neurons represents one possible explanation. The inclusion of the *Ptbp1*-dependent splicing of proapoptotic gene *Bak1* is essential for neuronal and animal survival^31^. Knockdown of residual PTBP1 expression in neurons may therefore further promote neuronal survival. Another possibility is unexpected beneficial off-target effects of reagents targeting *Ptbp1* in the previous reports. In either case, genetic methods should be used to validate future reports of glia-to-neuron conversion.

## Supplemental figure legends

**Extended Data Figure 1.**
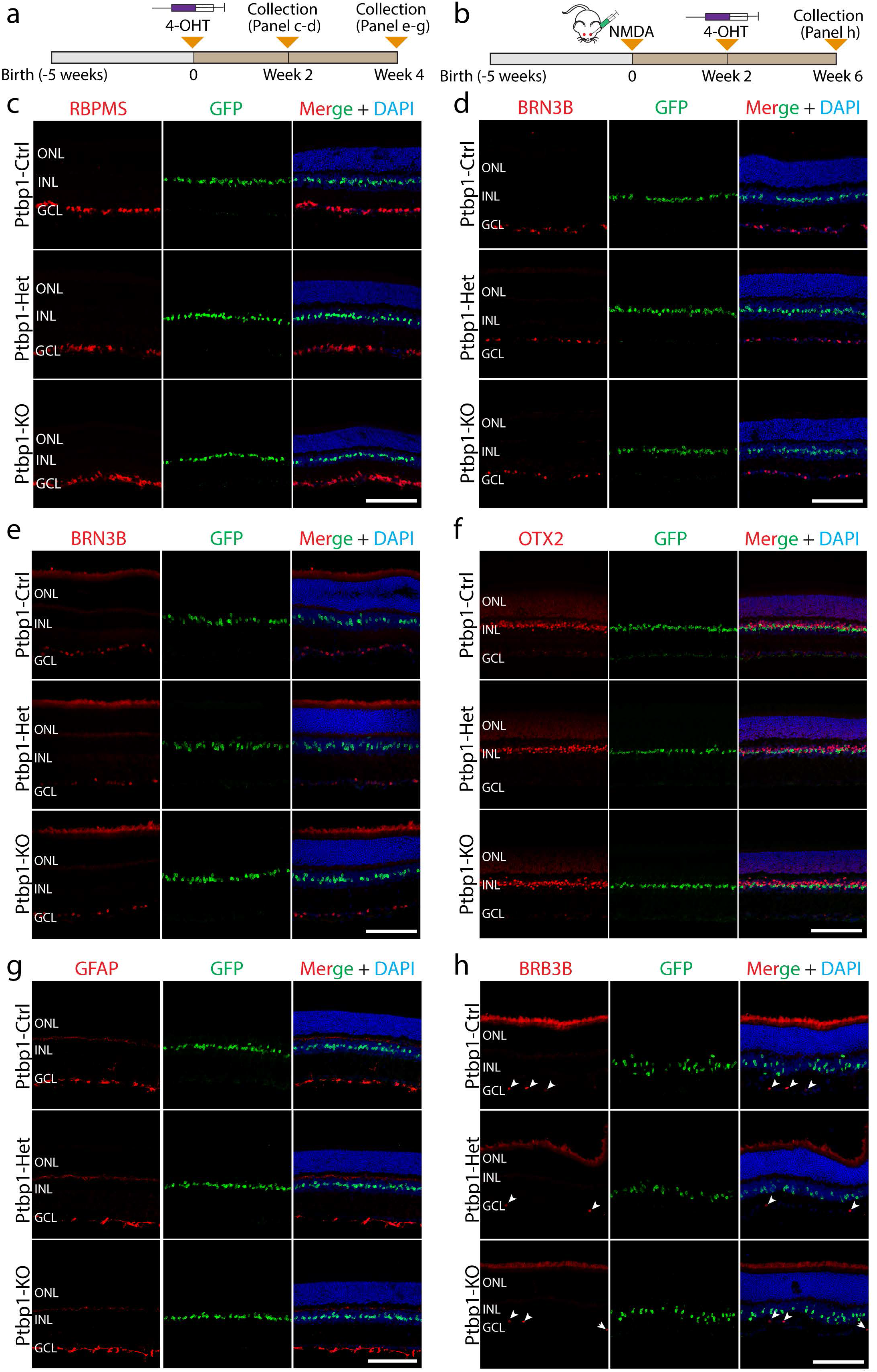
Immunohistochemical characterization of adult mouse retinas with Müller glia-specific *Ptbp1* deletion. **a, b**, A schematic diagram of the generation of specific *Ptbp1* deletion and lineage tracing of Müller glia and experimental timelines. **c, d**, Representative immunostaining for RBPMS (**c**) and BRN3B (**d**) expression in adult wildtype, heterozygous and homozygous *Ptbp1*-deleted mouse retinas after 2 weeks of 4-OHT i.p injection. No RBPMS/GFP-double positive or BRN3B/GFP-double positive was observed in any of the three genotypes (n >= 3 mice/genotype). **e, f, g**, Representative images of immunohistochemical analysis for BRN3B (**e**), OTX2 (**f**), and GFAP (**g**) expression in the retinas of the three genotypes after 4 weeks of 4-OHT i.p injection. No BRN3B/GFP-double positive or OTX2/GFP-double positive was observed in any of the three brain regions (n >= 3 mice/genotype). **h**, Efficient depletion of BRN3B-positive retinal ganglion cells in all three genotypes after 6 weeks of intravitreal NMDA injection. White arrowheads indicate remaining BRN3B-positive retinal ganglion cells that survived after intravitreal NMDA injection. ONL, outer nuclear layer; INL, inner nuclear layer; GCL, ganglion cell layer. Scale bars = 100 µm.

**Extended Data Figure 2.**
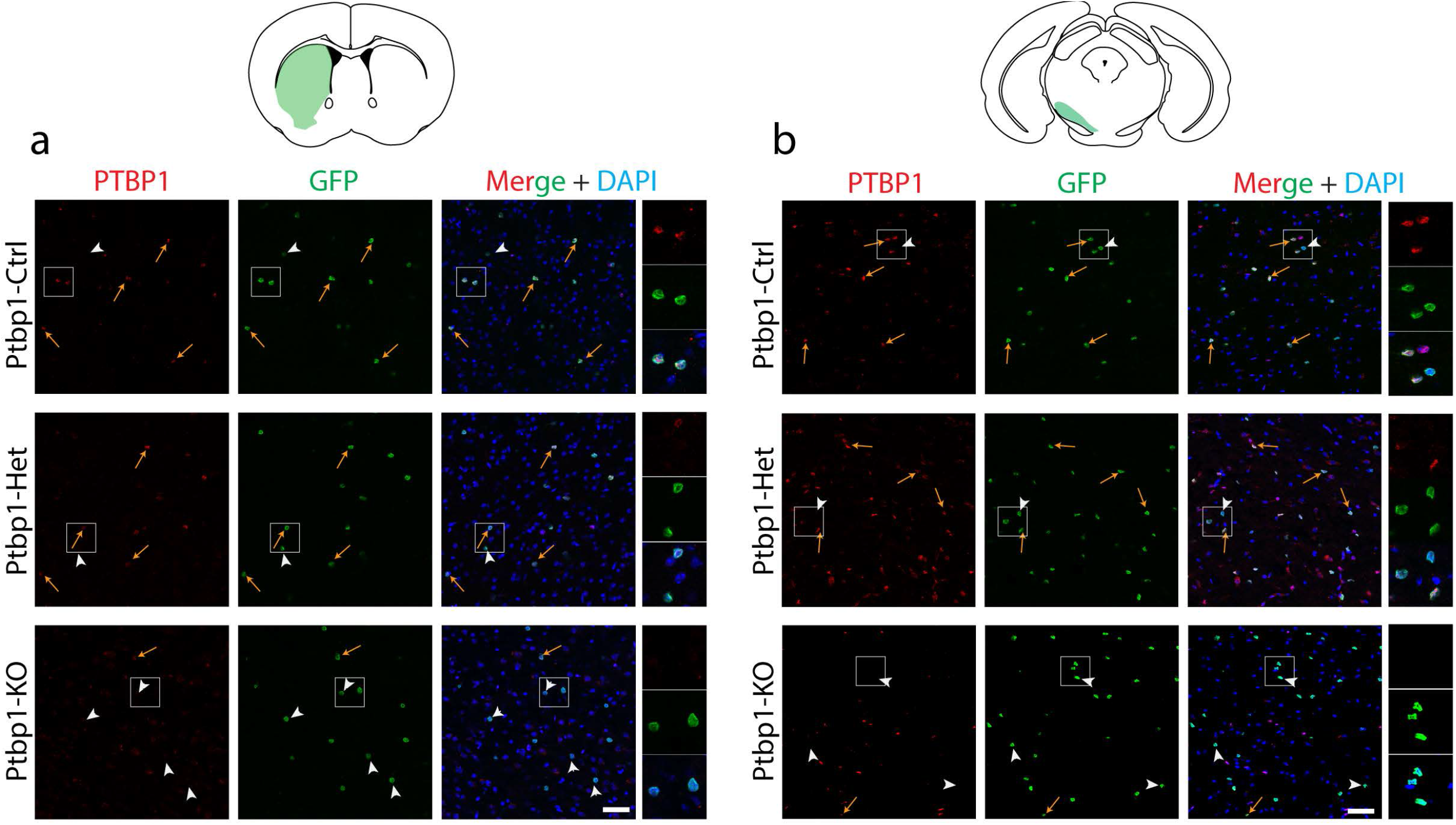
*Ptbp1* immunohistochemical analysis in adult mouse brain after astrocyte-specific deletion of *Ptbp1*. **a, b**, Representative PTBP1 and GFP immunostaining images of the striatum (**a**), substantia nigra (**b**) at 8 weeks after 4-OHT i.p injection (n >= 3 mice/genotype). Yellow arrows indicate PTBP1/GFP-double positive cells. White arrowheads indicate PTBP1-negative/GFP-positive cells. Scale bars = 50 µm.

**Extended Data Figure 3.**
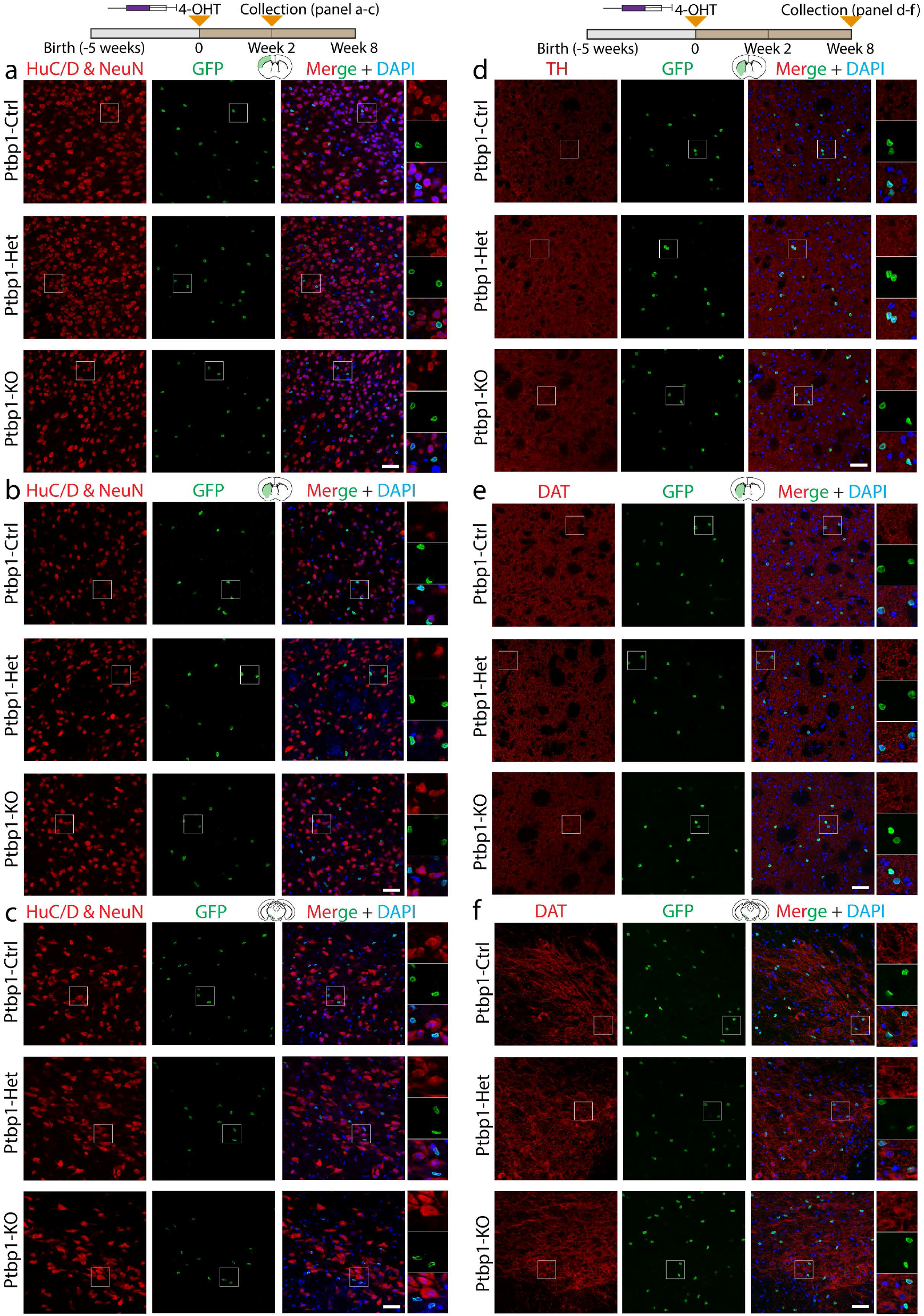
Immunohistochemical analysis of adult mouse brain after astrocyte-specific deletion of *Ptbp1*. **a-c**, Representative HuC/D;NeuN and GFP immunostaining images of the cortex (**a**), striatum (**b**), substantia nigra (**c**) at 2 weeks after 4-OHT i.p injection. No HuCD;NeuN/GFP-double positive cells were observed in any of the three brain regions across all 3 genotypes. **d**, Representative immunostaining images of TH and GFP in the striatum at 8 weeks after 4-OHT i.p injection. No TH/GFP-double positive cells were observed in the striatum in all 3 genotypes. Note that TH immunostaining signals are only present in non-cell bodies in the striatum. **e, f**, Representative DAT and GFP immunostaining images of the striatum (**e**), substantia nigra (**f**) at 8 weeks after 4-OHT i.p injection. No DAT/GFP-double positive cells were observed in any of the two brain regions across all 3 genotypes. Note that DAT immunostaining signals are only present in non-cell bodies in the striatum Scale bars = 50 µm.

**Extended Data Figure 4.**
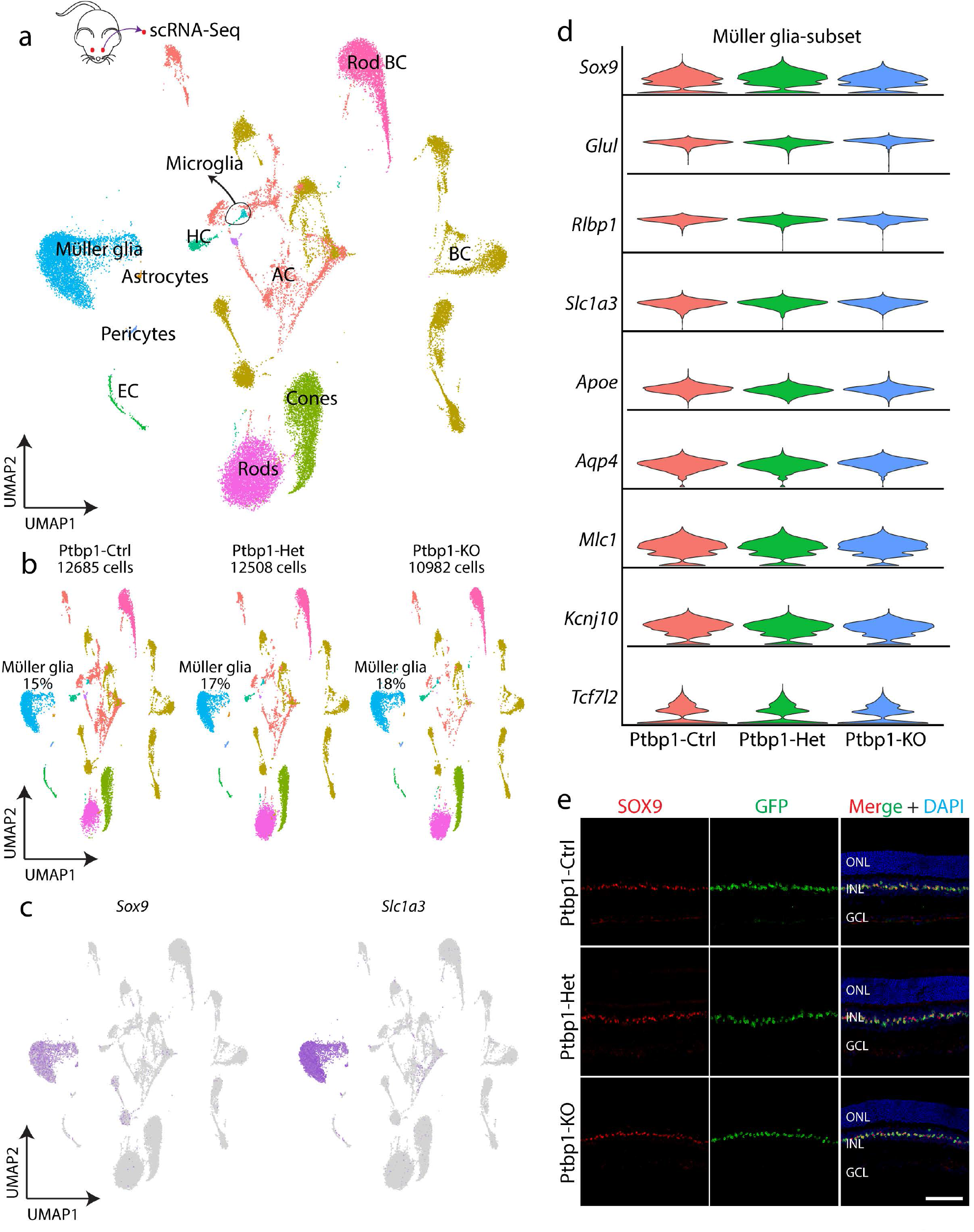
ScRNA-analysis of mouse retinas after Müller glia-specific *Ptbp1* deletion. **a, b**, UMAP plots showing scRNA-Seq clusters of all 3 combined genotypes (**a**) and separately (**b**), and the percentage of Müller glia in each scRNA-Seq dataset. **c**, Gene plots of representative Müller glia markers, *Sox9* and *Slc1a3*. **d**, Violin plots showing no significant change in the expression of Müller glia markers across 3 genotypes. **e**, Representative SOX9 and GFP immunostaining images of the retina from 3 genotypes at 4 weeks after 4-OHT i.p injection (n >= 4 mice/genotype). ONL, outer nuclear layer; INL, inner nuclear layer; GCL, ganglion cell layer. Scale bars = 100 µm.

**Extended Data Figure 5.**
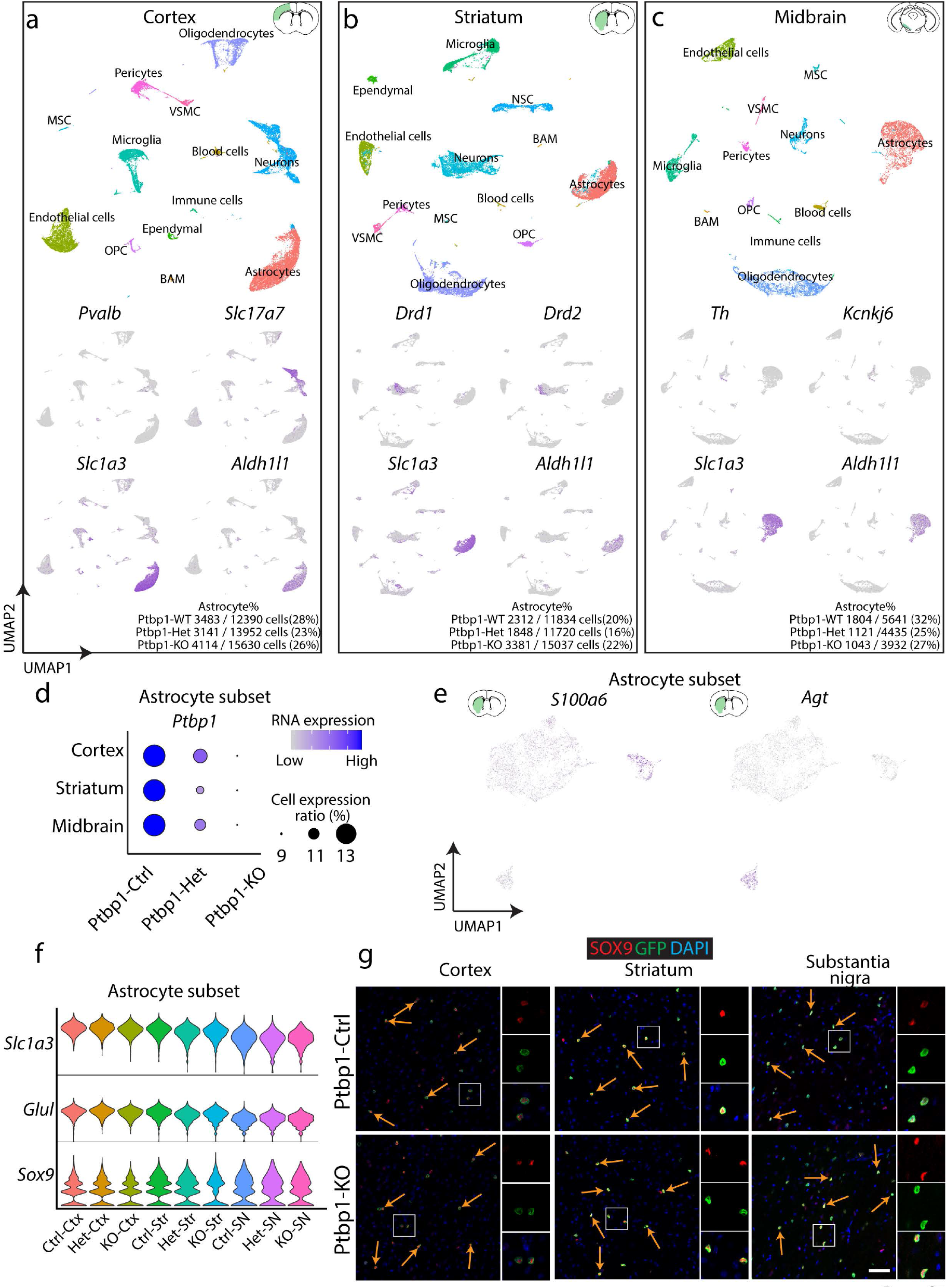
ScRNA-analysis of mouse brain after astrocyte-specific *Ptbp1* deletion. **a-c**, UMAP plots showing scRNA-Seq clusters of all 3 genotypes (top) and gene plots of representative neuronal markers and astrocyte markers in the cortex (**a**), striatum (**b**), and substantia nigra (**c**), and the percentage of astrocytes in each scRNA-Seq dataset across 3 genotypes. **d**, Dot plots showing *Ptbp1* expression in subsetted astrocytes in the 3 brain regions across 3 genotypes. **e**, Gene plot showing 2 sub-clusters in striatum astrocytes. **f**, Violin plots showing expression of astrocyte-enriched genes in subsetted astrocytes across 3 genotypes. **g**, Representative SOX9 and GFP immunostaining images of the cortex, striatum, and substantia nigra after at 8 weeks 4-OHT i.p injection. Scale bars = 50 µm.

## Supplemental Tables

**Supplemental Table 1**: Differential gene expression as measured by scRNA-Seq in retinal Müller glia across *Ptbp1* genotypes.

**Supplemental Table 2**: Differential gene expression as measured by scRNA-Seq in cortical astrocytes across *Ptbp1* genotypes.

**Supplemental Table 3**: Differential gene expression as measured by scRNA-Seq in striatal astrocytes across *Ptbp1* genotypes.

**Supplemental Table 4**: Differential gene expression as measured by scRNA-Seq in substantia nigra astrocytes across *Ptbp1* genotypes.

## Materials and Methods

### Mice

All experimental procedures were pre-approved by the Institutional Animal Care and Use Committee (IACUC) of the Johns Hopkins University School of Medicine and/or the Cleveland Clinic. *Glast*^*CreERT2*^ and *Sun1-GFP*^*lox/lox*^ transgenic mice were provided by Dr. Jeremy Nathans^23,25^. *Aldh1l1*^*CreERT2*^ transgenic mouse line, which allows for specific inducible Cre-recombination in astrocytes after TAM injection^26^, was purchased from JAX (JAX#029655). *Ptbp1*^*lox/lox*^ mice carrying loxP sites that flank the promoter and 1^st^ exon of *Ptbp1* were generated as described previously^32^. To induce specific Cre activation in adult Müller and astrocytes, 4 consecutive doses of 4-Hydroxytamoxifen (4-OHT) intraperitoneal (i.p) injection (30 mg/kg) were performed in adult wildtype, heterozygous and homozygous mice at ∼5 weeks old. Mice were sacrificed at the indicated time for analysis. All procedures were approved by the

### Intravitreal NMDA injection

We followed a previously described protocol^12^. Briefly, adult mice were anesthetized with 4% isoflurane inhalation. 1.5 μl of 200 mM NMDA in PBS was injected intravenously into the retinas. 4 consecutive doses of 4-OHT i.p injection (30 mg/kg) were performed at 2 weeks after NMDA injection. Mice were sacrificed, and retinas were collected at 4 weeks after 4-OHT injection.

### Immunohistochemical and imaging analysis

Retinal collection and immunohistochemical analysis were performed as described previously^24^. Briefly, mice were anesthetized by CO_2_, and eye globes were fixed in 4% paraformaldehyde in PBS for 4 hr at room temperature. Retinas were dissected and placed into 30% sucrose in PBS at 4°C overnight. Retinas were embedded in OCT, sectioned at 16 µm thickness, and stored at -20°C. Retinal sections were air-dried at 37°C for 20 min, washed 3 times x 5 mins with PBS, and incubated in a blocking buffer (0.4% TritonX-100, 10% horse serum in PBS) for 1 hr at room temperature. Primary antibodies were incubated in the blocking buffer at the indicated concentration overnight at 4°C. Primary antibodies used in this study were: rabbit anti-Ptbp1(1:200, Proteintech, #125821-AP), rabbit anti-Rbpms (1:200, Proteintech, #151871-AP), goat anti-Brn3b (1:200, Santa Cruz, #sc6026), rabbit anti-cone arrestin (1:200, Millipore Sigma, #AB15282), goat anti-Otx2 (1:200, R&D systems, #AF1979), rabbit anti-Gfap (1;300, Dako, #z0334), rabbit anti-Sox9 (1:200, Millipore Sigma, #AB5535), rabbit anti-GFP (1:400, Life technologies, #A6455) and chicken anti-GFP (1:400, Thermo Fisher Scientific, #A10262). Secondary antibodies were incubated at 1:400 dilution in the blocking buffer. Retinal sections were then incubated with DAPI, washed 4 times x 5 min in PBS, mounted with prolong gold mounting media (ThermoFisher Scientific, #P36935), air-dried, and stored at 4°C.

Brain collection and immunohistochemical analysis were performed as described previously^24,33,34^. Briefly, mice were anesthetized by avertin and cardially perfused with 4% paraformaldehyde. Brains were then dissected and post-fixed in 4% paraformaldehyde for 12 hr at room temperature, and then placed into 30% sucrose in PBS at 4°C overnight. Brains were sectioned at 30 µm thickness and processed for immunohistochemistry as described above. Additional primary antibodies used in this study for the brain were: mouse anti-HuC/D (1:200, Thermo Fisher Scientific, #A21271), mouse anti-NeuN (1:200, Millipore Sigma, #MAB377), rat anti-Dopamine Transporter Antibody (DAT, 1:200, Millipore Sigma, #MAB369), rabbit anti-Tyrosine Hydroxylase (TH, 1:200, Millipore Sigma, #AB152).

Images were acquired using Zeiss LSM700 confocal microscope at > 6 random regions for each retina or brain. Images were processed using ImageJ. All data were analyzed statistically by one-way ANOVA with a post hoc t-test for multiple comparisons using Excel. In all tests, values of *p < 0*.*05* were considered to indicate significance.

### Electroretinogram analysis

Mouse ERGs were recorded using a published procedure^35^. After overnight dark adaptation, mice were anesthetized using intraperitoneal injection of ketamine (80 mg/kg) and xylazine (16 mg/kg) and placed on a temperature-controlled heating pad. The pupils were dilated with eye drops (1% tropicamide, 1% cyclopentolate, 2.5% phenylephrine HCl). Strobe flash stimuli were presented within a UTAS Bigshot system (LKC Technologies) first in darkness (−3.6 to 2.1 log cd s/m^2^) and then superimposed upon a steady 20 cd/m^2^ achromatic background (–0.8 to 1.9 log cd s/m^2^). ERGs were recorded (0.3 – 1,500 Hz) using a stainless-steel wire electrode contacting the anesthetized (proparacaine HCl) corneal surface through a layer of 1% methylcellulose. The a-wave amplitude was measured at 8 msec after flash onset relative to the pre-stimulus baseline. The b-wave amplitude was measured from the a-wave trough to the peak of the b-wave, or from the pre-stimulus baseline if the a-wave was not detectable.

### Slice electrophysiology of cortical astrocytes

The acute brain slices were generated as previously described^36,37^. Ptbp1-Het or Ptbp1-KO (P54 to P73, male) were anesthetized with isoflurane and decapitated, and the brains were rapidly removed and chilled in ice-cold sucrose solution containing 76 mM NaCl, 25 mM NaHCO_3_, 25 mM glucose, 75 mM sucrose, 2.5 mM KCl, 1.25 mM NaH_2_PO_4_, 0.5 mM CaCl2, and 7 mM MgSO_4_ (pH 7.3). Acute brain slices (300 µm) including the motor cortex were prepared using a vibratome (VT 1200 s, Leica) and transferred to warm (32°C to 35°C) sucrose solution for 30 min for recovery. The slices were transferred to warm (32°C to 34°C) aCSF composed of 125 mM NaCl, 26 mM NaHCO_3_, 2.5 mM KCl, 1.25 mM NaH_2_PO_4_, 1 mM MgSO_4_, 20 mM glucose, 2 mM CaCl_2_, 0.4 mM ascorbic acid, 2 mM pyruvic acid, and 4 mM l-(+)-lactic acid (pH 7.3, 315 mOsm) and allowed to cool to room temperature. All solutions were continuously bubbled with 95% O_2_/5% CO_2_.

For whole-cell patch-clamp recordings, slices were transferred to a submersion chamber on an upright microscope [Zeiss Axio Examiner, objective lens: 5×, 0.16 numerical aperture (NA) and 40×, 1.0 NA] fitted for infrared differential interference contrast (IR-DIC) and fluorescence microscopy. Slices were continuously superfused (2 to 4 ml/min) with warm, oxygenated aCSF (32°C to 34°C). Sun1-GFP(+) cells in the layer 2/3 of the motor cortex were identified under a digital camera (Sensicam QE, Cooke) using transmitted light and green fluorescence. For the whole-cell patch-clamp recordings, borosilicate glass pipettes (2 to 4 MΩ) were filled with an internal solution containing 2.7 mM KCl, 120 mM KMeSO_4_, 9 mM HEPES, 0.18 mM EGTA, 4 mM Mg-adenosine 5′-triphosphate, 0.3 mM Na-guanosine 5′-triphosphate, and 20 mM phosphocreatine(Na) (pH 7.3, 295 mOsm). Whole-cell patch-clamp recordings were conducted through a MultiClamp 700B amplifier (Molecular Devices) and an ITC-18 (InstruTECH), which were controlled by customized routines written in Igor Pro (WaveMetrics).

To determine the action potential firing capacity of the recorded cells, a series of depolarizing current steps (1-s long, 0 to 4000 pA) was given to the cells, and their voltage responses were recorded. During the current step test, cells were held at their resting membrane potentials, and different intensities of currents were tested until the cell membrane potential went over 0 mV without firing action potentials or the membrane potential did not hold from 4000 pA current injections. As a control, three Sun1-GFP(-) neurons from two Ptbp1-KO mice were targeted and their voltage responses to a series of depolarizing current steps (1-s long, 0 to 300 pA) were recorded by using the same internal and external solutions for comparing action potential firing properties with Sun1-GFP(+) cells from the same mice. All signals were low-pass filtered at 10 kHz and sampled at 20 kHz. Electrophysiology data was analyzed in Igor Pro (WaveMetrics).

### Cell dissociation and scRNA-Seq

Retinal cell dissociation was performed as described previously^24^. Briefly, one female mouse per genotype was euthanized by CO_2_, and eye globes were removed and placed in ice-cold PBS. Retinas were dissected, cells were dissociated using Papain Dissociation System (LK003150, Worthington). Dissociated cells were resuspended in a buffer containing 9.8 ml Hibernate A, 200 μl B27, 20 μl GlutaMAX and 0.5 U/μl RNAse inhibitor. Cells were filtered through a 50 μm filter. Cell count and viability were determined by using 0.4% Trypan blue.

Brain cell dissociation was performed as described previously^38,39^ using one female mouse per genotype. Mouse brain matrix (0.5 mm) was used to slice brains into coronal sections, and relative brain regions (cerebral cortex, striatum, midbrain) were dissected into Hibernate-A media with a 2% B-27 and GlutaMAX supplement (0.5 mM final). Tissues were dissociated in papain (Worthington).

Cells were then loaded into the 10x Genomics Chromium Single Cell System (10x Genomics) and libraries were generated using v3.1 chemistry following the manufacturer’s instructions. Libraries were sequenced on the Illumina NovaSeq platform (500 million reads per library).

### ScRNA-Seq data analysis

Sequencing data were first processed through the Cell Ranger (v6.0.1, 10x Genomics) with default parameters, aligned to the mm10 genome (refdata-gex-mm10-2020-A), and matrix files were used for subsequent bioinformatic analysis.

Matrix data were further processed using Seurat 3.4 version^40^. Low quality cells with < 500 genes, < 2000 UMI and > 30% mitochondrial genes were removed. Datasets were normalized using Seurat ‘scTransform’ function. Cells were clustered and visualized using UMAP. Retinal Müller glia (*Slc1a3*^+^ & *Rlbp1*^+^) and brain astrocytes (*Slc1a3*^+^ & *Aldh1l1*^+^) clusters were subsetted for further analysis to compare differential gene expression across genotypes. Differential gene expression between different genotypes was calculated using the Seurat ‘*FindMarker’* function.

## Acknowledgments

We thank F. Zhou, A. Fischer, J. Ling, Cheng Qian, and W. Yap for their comments on the manuscript. We thank the Single Cell & Transcriptomics Core (Johns Hopkins) for sequencing of scRNA-Seq libraries. This work was supported by the NIH National Eye Institute grants R01EY020560 and U01EY027267 to SB, R24EY027283 to SB and NSP, Core Grant P30EY025585 to NSP, and the Maryland Stem Cell Research Fund (2019-MSCRFF-5124) to DWK. SB is supported by a Stein Innovation Award from Research to Prevent Blindness.

## Author contributions

TH, DWK, and SB conceived and supervised the study. TH generated and analyzed immunohistochemistry and scRNA-Seq data from the retina, assisted by HA, NAP, and DWK. DWK generated and analyzed immunohistochemistry and scRNA-Seq data from the brain, assisted by TH and HA. MO and SZ provided reagents. JK performed slice electrophysiology of cortical astrocytes. MY and NSP conducted ERG analysis. TH, DWK, and SB drafted the manuscript. All authors contributed to the writing of the manuscript.

## Statement of interests

SB is a co-founder of, and shareholder in, CDI Labs LLC, and is a consultant to Third Rock Ventures, LLC.

## Data availability

All scRNA-Seq data have been deposited to GEO as GSE184933.

## Notes

### Summary of Updates

ERG data added. Figure 1, manuscript text, and methods modified.

